# Endothelial reprogramming by disturbed flow revealed by single-cell RNA and chromatin accessibility study

**DOI:** 10.1101/2020.07.15.205427

**Authors:** Aitor Andueza, Sandeep Kumar, Juyoung Kim, Dong-Won Kang, Hope L Mumme, Julian I Perez, Hanjoong Jo

## Abstract

Disturbed flow *(d-flow)* induces atherosclerosis by regulating gene expression in endothelial cells (ECs). For further mechanistic understanding, we carried out a single-cell RNA sequencing (scRNAseq) and scATACseq study using endothelial-enriched single-cells from the left- and right carotid artery exposed to d-flow (LCA) and stable-flow (*s-flow* in RCA) using the mouse partial carotid ligation (PCL) model. We found 8 EC clusters along with immune cells, fibroblasts, and smooth muscle cells. Analyses of marker genes, pathways, and pseudo-time revealed that ECs are highly heterogeneous and plastic. *D-flow* induced a dramatic transition of ECs from atheroprotective phenotypes to pro-inflammatory, mesenchymal (EndMT), hematopoietic stem cells, endothelial stem/progenitor cells, and an unexpected immune cell-like (EndICLT) phenotypes. While confirming KLF4/KLF2 as *s-flow*-sensitive transcription factor binding site, we also found those sensitive to d-flow (RELA, AP1, STAT1, and TEAD1). D-flow reprograms ECs from atheroprotective to pro-atherogenic phenotypes including EndMT and potentially EndICLT.

## INTRODUCTION

Atherosclerosis is the major underlying cause of myocardial infarction, ischemic stroke, and peripheral arterial disease (PAD), which represent a leading cause of death worldwide (Herrington et al., 2016). Atherosclerosis is a chronic inflammatory disease and occurs preferentially in arterial regions exposed to disturbed blood flow *(d-flow)*, while those exposed to stable flow *(s-flow)* are protected (Chiu and Chien, 2011; Kwak et al., 2014; Tarbell et al., 2014). Flow is recognized by mechanosensors in endothelial cells (ECs), which in turn activate signaling pathways leading to regulation of gene expression, endothelial function, and atherogenic pathways (Simmons et al., 2016). *D-flow* induces crucial pro-atherogenic pathways in ECs including endothelial inflammation and dysfunction, permeability dysfunction, thrombosis, and endothelial-to-mesenchymal transition (EndMT). In contrast, *s-flow* protects ECs from those proatherogenic pathways (Kumar et al., 2014; Mahmoud et al., 2017; Nigro et al., 2011).

To understand how flow regulates expression of endothelial genes at the transcript and epigenetic levels *in vivo*, we and others used a bulk RNAs or DNAs obtained from the pooled ECs exposed to either *s-flow* or *d-flow* using the mouse partial carotid ligation (PCL) model or pig arteries (Dunn et al., 2014; Jiang et al., 2015; Ni et al., 2010; Son et al., 2013). We developed the PCL model and showed that *d-flow* rapidly induces, whereas *s-flow* prevents, robust atherosclerosis development within two weeks in hypercholesterolemic mice (Kumar et al., 2017; Nam et al., 2010; Son et al., 2013). The PCL model involves the ligation of 3 of 4 distal branches of the left common carotid artery (LCA) to induce *d-flow*, while using the contralateral right common carotid artery (RCA) that continues to be exposed to *s-flow* as an internal control. We further developed a lumen flushing method that enabled us to obtain endothelial-enriched RNAs and DNAs from the LCAs and RCAs following the PCL. These pooled bulk RNAs were then analyzed by mRNA microarray, microRNA microarray, and RNA sequencing to identify mRNA transcriptomes and microRNAs regulated by *flow* in ECs. These studies led to the discovery of numerous flow-sensitive genes and microRNAs that have been characterized and studied in detail for their roles in endothelial biology and atherosclerosis. We were also able to obtain bulk DNA samples from the LCAs and RCAs and analyzed by a reduce-representation bisulfite sequencing (RRBS) method (Dunn et al., 2014). This study showed that *d-flow* and *s-flow* differentially regulate epigenomic DNA methylation profiles in ECs, identifying many gene loci that were regulated by flow.

While these transcriptome and epigenomic methylome studies using the bulk RNA and DNA samples clearly identified the differential effects of *d-flow* and *s-flow* on ECs *in vivo*, the interpretation of these studies had several limitations. While our bulk RNA and DNA preparations were highly enriched with ECs from the luminal flushings, it was unavoidable that they contained some other cell types present in the intima especially in the LCAs following the PCL surgery. For example, we observed an increase in the expression of many genes associated with immune cells and mesenchymal cells in the LCA samples as early as 12 h following PCL; however, we could not ascertain whether those changes were due to non-ECs infiltrating to the intima or were indeed induced by *d-flow* in ECs.

Recent development in single-cell RNA sequencing (scRNAseq) and single-cell Assay for Transposase Accessible Chromatin sequencing (scATACseq) methods has enabled the transcriptomic and epigenomic analyses of a large number of cells at single-cell resolution at once. While scRNAseq study shows an abundance of each gene transcript in individual cells and provides insights into gene transcript expression profile, scATACseq analysis reveals changes in the chromatin accessibility profile allowing insights into epigenomic regulation of gene expression genome-wide. Further, scATACseq analysis reveals several layers of gene regulation, such as genome-wide identification of cis-regulatory elements including enhancers, nucleosome positions, and transcription factor (TF) binding sites. Since gene expression is regulated both at the transcriptional and epigenomic levels, the unprecedented power of carrying out concomitant and integrated analyses of scRNAseq and scATACseq data are increasingly evident. These new powerful tools have revealed the heterogeneity of cell types in the liver (Zhao et al., 2020), lungs (Domingo-Gonzalez et al., 2020), heart (Bykov et al., 2020), immune cells (Villani et al., 2017), and ECs (Kalluri et al., 2019; Kalucka et al., 2020) under various physiological and pathological conditions.

Here, we performed a scRNAseq and a scATACseq assay using the endothelial-enriched single cells and nuclei obtained from the mouse carotid arterial lumens, following the PCL to determine the differential effect of *d-flow* and *s-flow* on genome- and epigenome-wide regulation of gene transcripts and chromatin accessibility profiles. Individual analyses of scRNAseq and scATACseq data as well as an integrated analysis of both data sets revealed that ECs are heterogenous in the carotid artery even under *s-flow* condition. *D-flow* dramatically altered endothelial transcriptome and epigenomic profiles, reprogramming them into inflammatory, mesenchymal, stem/progenitor-like cells, hematopoietic cells, and immune cell-like phenotypes. In addition, we also identified a *s-flow* and *d-flow* dependent enrichment of TF binding sites and cis-regulatory elements.

## RESULTS

### scRNAseq and scATACseq analyses using the mouse PCL model

To determine the genome-wide effect of *s-flow* and *d-flow* on gene transcript expression and chromatin accessibility profiles in ECs at a single-cell resolution *in vivo*, we carried out a scRNAseq and scATACseq using the mouse PCL model (Figure S1). C57BL/6 mice were partially ligated to induce *d-flow* in the LCA using the contralateral RCA exposed to *s-flow* as controls. Two days (2D) and two weeks (2W) post partial ligation, RCAs (2D-R and 2W-R) and LCAs (2D-L and 2W-L) were luminally digested with collagenase to obtain single-cells and single-nuclei, which were used for scRNAseq and scATACseq, respectively.

### scRNAseq analysis

Single cells from 10 RCAs and LCAs were prepared, barcoded, and sequenced. The number of single-cells used for scRNAseq were total of 9,709 cells pooled from 1,867 (2D-R), 2,119 (2D-L), 1,263 (2W-R), and 4,460 (2W-L) (Table S1). The mean reads per cell ranged from 34,142 to 88,631 and the genes expressed per cell were 2,770 to 3,537.

scRNAseq data analysis was performed using Seurat v3.6. As a quality control, we eliminated cells that expressed gene counts >7,600 or <200, and those expressing >10% mitochondrial unique molecular identifier (UMI) counts to remove doublets and damaged cells during the sample preparation. Following normalization, an unsupervised graph-based clustering partitioned the cells into groups as visualized by the Uniform Manifold Approximation and Projection (UMAP) plot (Figure 1A). Our UMAP analysis identified 16 unique cell clusters distributed in a flow- and time-dependent manner (Figure 1A and B). Each cell cluster was identified using well-established marker genes (Figure 1C) (Cochain et al., 2018; Kalluri et al., 2019; Roberts et al., 2020; Winkels et al., 2018). We found 8 endothelial cell clusters (E1-E8), vascular smooth muscle cells (SMCs), fibroblasts (Fibro), 4 monocytes/macrophages (Mϕ1-4), dendritic cells (DCs) and T-cells. Endothelial cell clusters were identified based on the expression of *Pecam1, Icam2, Cdh5*, and *Tie1. SMC*-specific markers were *Cnn1, Myl9*, and *Speg.* Likewise, *Medag, Dpep1, and Lum* were used as markers for Fibro; *C1qa, C1qb, C1qc*, and *C5ar1* for Mϕs; *Ccr7* and *Mmp25* for DCs; and *Itk* for T-cells (Figure 1C).

**Figure 1.**
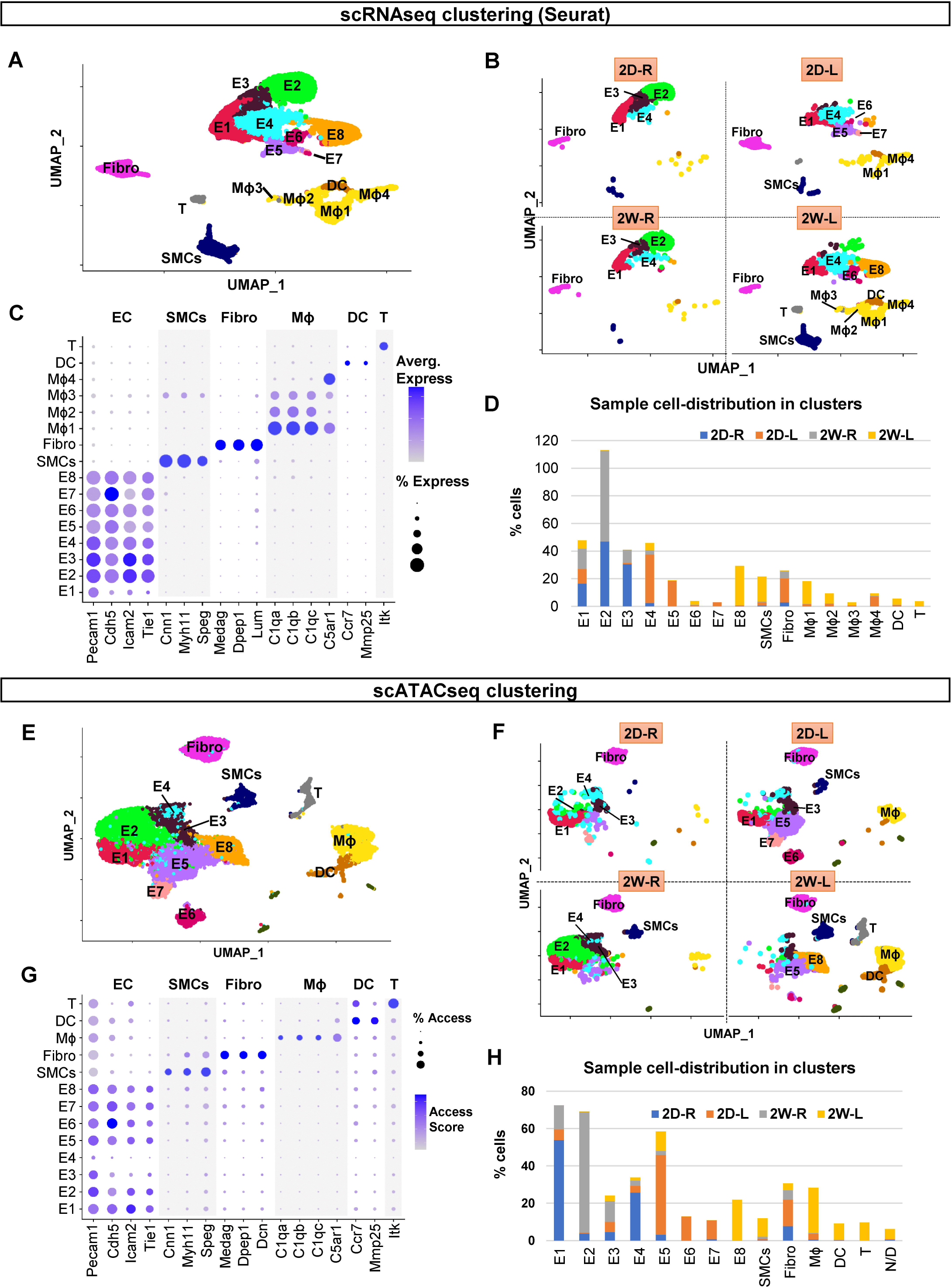
scRNAseq and scATACseq clustering and cell identification. **(A and B)** scRNAseq data plotted on a single UMAP representing 2D-R, 2D-L, 2W-R and 2W-L. **(A)** shows a UMAP representing all 4 samples while **(B)** represents individual UMAP plots for each condition. Major cell populations include ECs (E1-8), SMCs, Fibro, Mϕs, DCs, and T-cells. **(C)** Dot plot shows the specific marker genes used to classify each cell clusters. The percentage of cells expressing respective transcripts correlate and their expression intensity are depicted by the dot size color intensity, respectively. **(D)** Graph shows the cell numbers in each cell cluster for 4 different experimental conditions. **(E and F)** scATACseq data plotted on a single UMAP representing 2D-R, 2D-L, 2W-R and 2W-L to identify major cell populations. **(E)** shows a single UMAP to identify major cell populations while **(F)** represents individual UMAP plots for each condition. **(G)** Dot plot shows the specific marker genes used to classify each cell clusters. The percentage of cells with open accessibility and the accessibility scores are depicted by the dot size and color intensity, respectively. **(H)** Graph shows the cell numbers in each cell cluster for 4 different experimental conditions.

The unique flow- and time-dependent cell cluster distribution pattern shown in Figure 1B was further quantified in Figure 1D. E1 cluster consisted of ECs present in all 4 conditions (2D-R, 2D-L, 2W-R and 2W-L). E1-E4 consisted of ECs exposed to *s-flow* conditions (2D-R and 2W-R). The majority of E5 and E7 clusters consisted of ECs exposed to acute *d-flow* (2D-L). Interestingly, E6 and E8 exclusively consisted of ECs exposed to chronic *d-flow* (2W-L). These results suggest that *d-flow* induced a phenotypic shift in a time-dependent manner. However, there was no discernable difference between the 2D-R and 2W-R samples in overall cell clusters, suggesting that the cells exposed to s-flow conditions for 2 days or 2 weeks remained largely unchanged. Few SMCs were found in *s-flow* conditions (2D-R and 2W-R), while SMC numbers increased in acute *d-flow* conditions (2D-L) to 2.3%, which further increased by the chronic *d-flow* (2W-L) to 18.2% of the total cells. Fibroblasts were present in all four conditions with the most in the acute *d-flow* condition (17% at 2D-L) (Figure 1D).

We found four Mϕ clusters, which showed dramatic flow- and time-dependent changes. In the *s-flow* conditions (2D-R and 2W-R), Mϕs were rare. However, acute *d-flow* increased the number of Mϕs to 10.3% (2D-L), which further increased to 28.4% in chronic d-flow (2W-L). This finding is consistent with the known effects of *d-flow* on immune cell accumulation (Alberts-Grill et al., 2012). Moreover, we found that Mϕ4 was the major population in the acute *d-flow* (2D-L), whereas Mϕ1 and Mϕ2 were the major populations found in the chronic *d-flow* (2W-L) group. This suggests a flow- and time-dependent phenotypic changes of these Mϕ populations. Like Mϕs, few DCs and T-cells were present in the *s-flow* conditions, while they increased by chronic *d-flow* (2W-L) to 4.6% and 3.7%, respectively.

The scRNAseq analysis performed above was independently validated by using another software, Partek Flow® which produced nearly identical results (Figure S2).

### scATACseq analysis

For this study, single nuclei from 12 RCAs and LCAs were prepared, subjected to transposase treatment, barcoded, and sequenced. The number of single-nuclei used for sequencing were total of 18,324, which were pooled from 1,291 (2D-R), 5,351 (2D-L), 5,826 (2W-R), and 5,856 (2W-L), respectively (Table S2). The mean fragments per cell ranged from 19,690 to 24,459 and the total reads ranged from 304 to 372 million reads per group.

The sequence reads from the four samples, 2D-R, 2D-L, 2W-R and 2W-L were analyzed using R software packages. The UMAP analysis partitioned total nuclei populations to 13 cell clusters, which were distributed in flow- and time-dependent manner (Figure 1E and F). The scATACseq data were converted to a gene-activity index by Signac, and cell identities were determined using the same marker genes described in the scRNAseq study above. Among the 13 clusters, 8 were endothelial clusters (E1-E8), followed by SMCs, Fibro, Mϕ, DCs, T-cells, and non-defined (ND) (Figure 1G). Unlike the scRNAseq result, only one Mϕ cluster was identified.

The unique flow-and time-dependent cell cluster distribution pattern shown in Figure 1F was further quantified in Figure 1H. Like the scRNAseq analysis, E1-E4 clusters consisted of ECs exposed to acute and chronic *s-flow* conditions (2D-R and 2W-R). Most E5-E7 clusters consisted of ECs exposed to acute *d-flow* (2D-L). While E6 and E7 were unique to ECs present in the acute *d-flow* condition (2D-L), E8 cells were exclusively found in the chronic *d-flow* condition (2W-L). As observed in the scRNAseq, most SMCs came from the chronic *d-flow* condition (2W-L), while the fibroblasts were found in all four conditions. The majority of Mϕs were found in the chronic *d-flow* condition (2W-L), followed by the acute *d-flow* (2D-L), while few were found in the *s-flow* conditions. Similarly, DCs and T-cells were found mainly in chronic *d-flow* conditions. These results suggest that E2 represents unique *s-flow* phenotype, E6 and E7 represent the intermediate phenotype in response to acute *d-flow*, while E8 is a unique phenotype induced by chronic *d-flow.*

Taken together, these scRNAseq and scATACseq results showed that *d-flow* alters genome-wide changes in gene expression and chromatin accessibility, leading to a dramatic shift in the endothelial phenotypes in a time-dependent manner. It also shows heterogeneous endothelial populations even within the carotid artery under the s-flow conditions, and *d-flow* further increases the heterogeneity *in vivo*.

### Integration of scRNAseq and scATACseq

Thus far, we analyzed the scRNAseq and scATACseq data independently. While scATACseq is used to profile epigenomic chromatin accessibility indicating the potential gene activity in each cell, scRNAseq data shows the transcript abundance in each cell, which is a different but complementary aspect of gene regulation. We integrated the scRNAseq and scATACseq datasets to correlate and cross-validate gene expression profiles and chromatin accessibility landscape in the luminal cells in response to *flow*. The co-embedded UMAP plot showed that >80% total cells analyzed in the scRNAseq and scATACseq datasets overlapped each other, suggesting that the changes in chromatin accessibility and corresponding gene transcript expression in most cell clusters occurred in a concordant manner (Figure 2A). The individual UMAP plots for the scATACseq and scRNAseq data are shown with the respective cell cluster identities (Figure 2B and C). Inspection of the remaining non-overlapping cells showed that E6’s, E7’s and parts of E1’s in both datasets did not show overlap, indicating that chromatin accessibility changes in these ECs were not highly concordant with the corresponding gene transcription profiles. As shown in Figure 1, E6’s and E7’s were the intermediate and transient phenotypes induced by acute *d-flow*, while E1 cells were derived from all four conditions.

**Figure 2.**
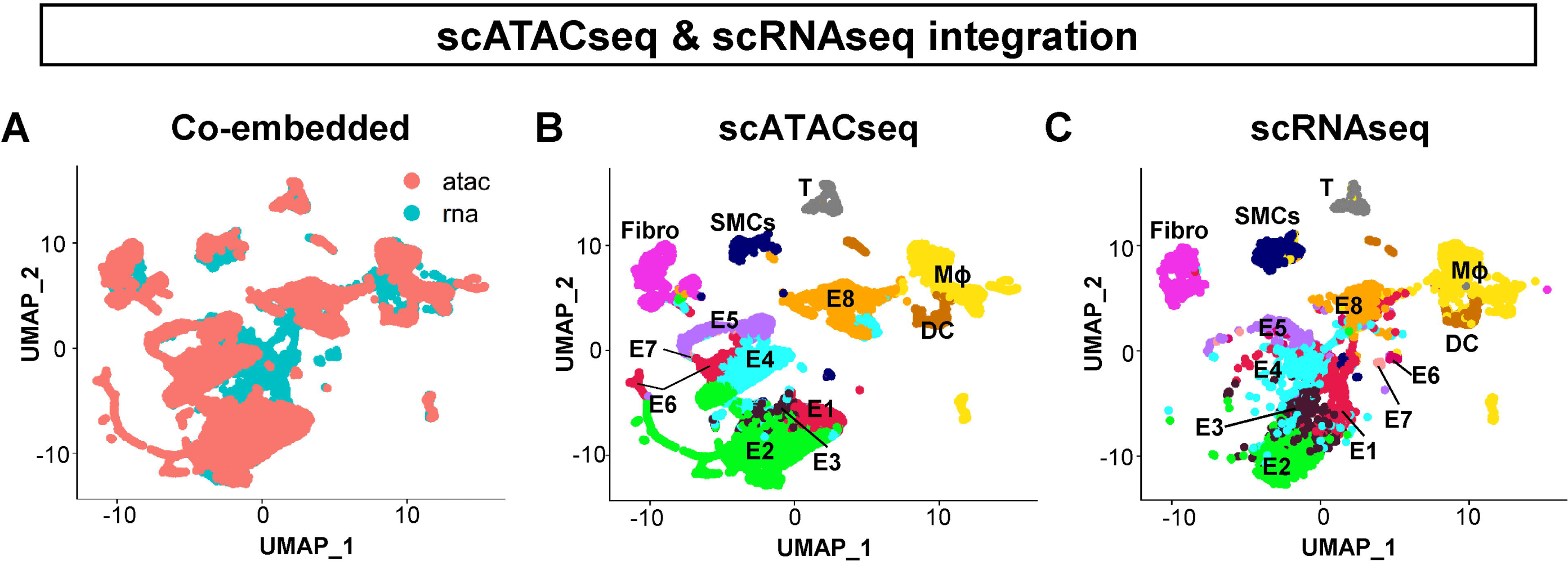
Integration of scRNAseq and scATACseq datasets. **(A)** shows a UMAP representation of the co-embedded scATACseq (Red) and scRNAseq (Cyan) datasets. Cells from two assays were labeled with two colors as indicated in the plot. **(B and C)** show the individual UMAP representation of the co-embedded scATACseq and scRNAseq datasets. Annotations used for the scRNAseq shown in Figure 1A were used and superimposed onto each cell cluster.

Taken together, these results obtained from two independent approaches demonstrate that our two datasets are highly concordant and cross-validate each other. This demonstrates the reproducible and robust effects of *d-flow* on genome-wide regulation of genes at the transcript and epigenomic chromatin accessibility levels in the aortic luminal cells at a single-cell level *in vivo*.

### D-flow induces pro-atherogenic pathways in ECs

Having established the high concordance between the scRNAseq and scATACseq data, we then determined the characteristics of each cell cluster by analyzing differential gene transcript expression patterns. For this, we carried out a differentially expressed feature analysis using the scRNAseq dataset and compared all the cell clusters with each other. Top 10 upregulated genes from each cell cluster were then used to generate a heatmap plot (Figure 3A). We found that E2 expressed the highest levels of two best-known mechanosensitive genes, *Klf2* and *Klf4*, followed by E3 (Figure S3). *Klf2* and *Klf4* expression were much lower in other EC clusters exposed to *d-flow* conditions (E5-E8). Interestingly, E1 and E4 clusters containing ECs from all four flow conditions showed a relatively low *Klf2* and *Klf4* expression, indicating that this heterogenous EC population is insensitive to flow. These results established E2 as a representative atheroprotective endothelial phenotype under *s-flow* condition. In contrast, three well-known *d-flow*-induced pro-atherogenic genes, *Ctgf, Serpine1, and Edn1* were highly expressed in ECs exposed to acute *d-flow* in E5. E8 cluster found in the chronic *d-flow* condition showed a well-known *d-flow*-induced gene *Thsp1.* In addition, E8 cells showed many genes that were also highly expressed in SMCs *(Acta2, Tagln)*, Fibro *(Dcn1), and* immune cells *(Cd74, H2-eb1, H2-aa, H2-ab1)*, indicating a potential for EndMT and acquisition of immune cell-like features by ECs under the chronic *d-flow* condition. Many of the top genes expressed in the non-endothelial clusters were well-known marker genes used for their identifications (Figure 1C). For example, SMCs showed high expression of *Acta2, Myh9, Myh11* and *Tgln*; Fibro expressed *Dcn*; Mϕs showed high expression of *C1qa, C1qb, C1qc* and *Ccl4*, respectively. Our attempt to compare the characteristics of the four Mϕ clusters did not reveal whether they represented a proinflammatory M1 or reparative M2 phenotypes (Alberts-Grill et al., 2012).

**Figure 3.**
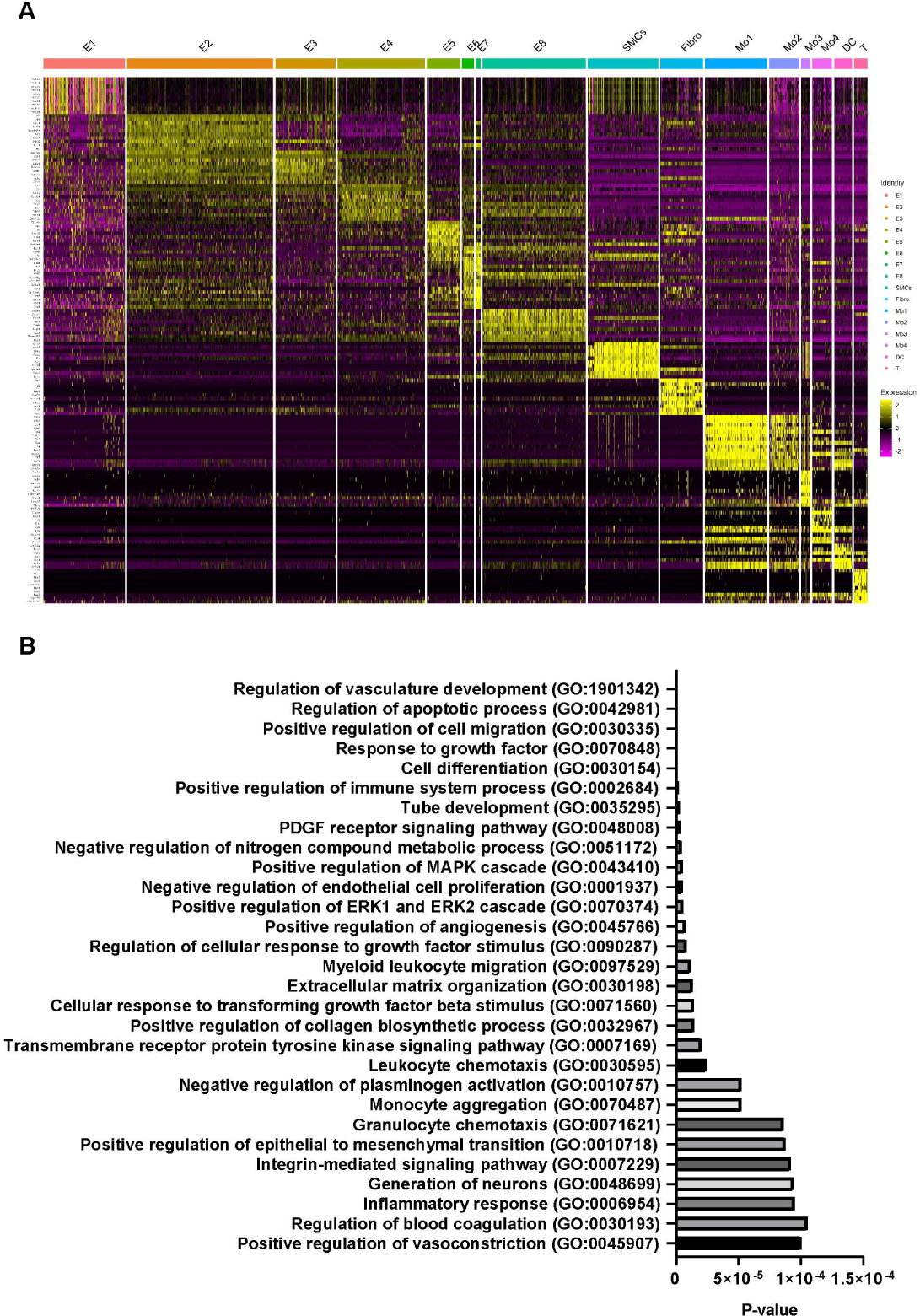
Differential gene expression and pathway analyses. **(A)** Heatmap represents the top 10 gene transcripts for each cell-cluster identified from the scRNAseq analysis. **(B)** Gene Ontology analysis was performed using the top 200 upregulated genes in E8 representing the *d-flow* phenotype in comparison to E2 representing the *s-flow* phenotype. X-axis shows the p-value.

To compare the biological status of the EC clusters exposed to *s-flow* and chronic *d-flow*, we selected respective E2’s and E8’s from both scRNAseq and scATACseq datasets, since they were clearly concordant and distinguishable from other endothelial clusters and represented the atheroprotective *s-flow* and pro-atherogenic *d-flow* conditions, respectively. The overexpressed genes found in both datasets in E8’s in comparison to E2’s were subjected to a Panther (http://www.pantherdb.org/) Gene Ontology (GO) analysis (Mi et al., 2017). As shown in Figure 3B, ECs exposed to chronic *d-flow* exhibited induction of many well-known biological processes associated with proatherogenic pathways (Simmons et al., 2016). These include vascular development, inflammation, apoptosis, angiogenesis, EndMT, TGF-β pathway and endothelial permeability (Daniel and van Buul, 2013; DePaola et al., 1999; Fan and Karino, 2010; Jenkins et al., 2013; Johnson and Nerem, 2007; Kutikhin et al., 2018; Rocha et al., 2018; Schnittler, 1998; Seebach et al., 2000; Ueno et al., 2000; Yamamoto et al., 2003). These results suggest that d-flow induces the transition of endothelial cells to pro-atherogenic phenotypes through by regulating gene transcripts and epigenomic chromatin accessibility profiles.

### Pseudo-time trajectory, chromatin accessibility, and gene expression analyses reveal flow-dependent endothelial reprogramming

The finding that ECs in E8 in response to chronic flow expressed the marker genes of mesenchymal and immune cells (Figure 3A) raised a question if *d-flow* induced endothelial reprogramming such as EndMT. To test this hypothesis, we conducted a trajectory analysis with Monocle 2 embedded in Partek Flow® software using the scRNAseq data to track dynamic differentiation status of each cell (Pliner et al., 2018). Our analysis clearly identified three major groups of differentiated cell types: 1) ECs, 2) immune cells (Mϕs, DC and T cells), and 3) SMCs and Fibros (Figure 4A). In addition, there were many other cells present along the trajectory paths towards each cell type, indicating their transitional de-differentiation status in response to *d-flow*. To better understand the trajectory results, we split the analysis by the four experimental conditions (Figure 4B). In the *s-flow* conditions (2D-R and 2W-R), most cells were well-differentiated with few cells in transition. In contrast, *d-flow* conditions induced many ECs to transitional states, potentially trans-differentiating towards SMCs and Fibro, indicating EndMT. Surprisingly, we also found many ECs transitioning towards immune cells in acute *d-flow* condition (2D-L), which further increased in the chronic *d-flow* (2W-L) condition. Further analysis using E8 cluster revealed that these ECs exposed to chronic *d-flow* transitioned toward immune cells (EndICLT), but did not achieve fully differentiated immune cell status (Figure 4C and Figure S4).

**Figure 4.**
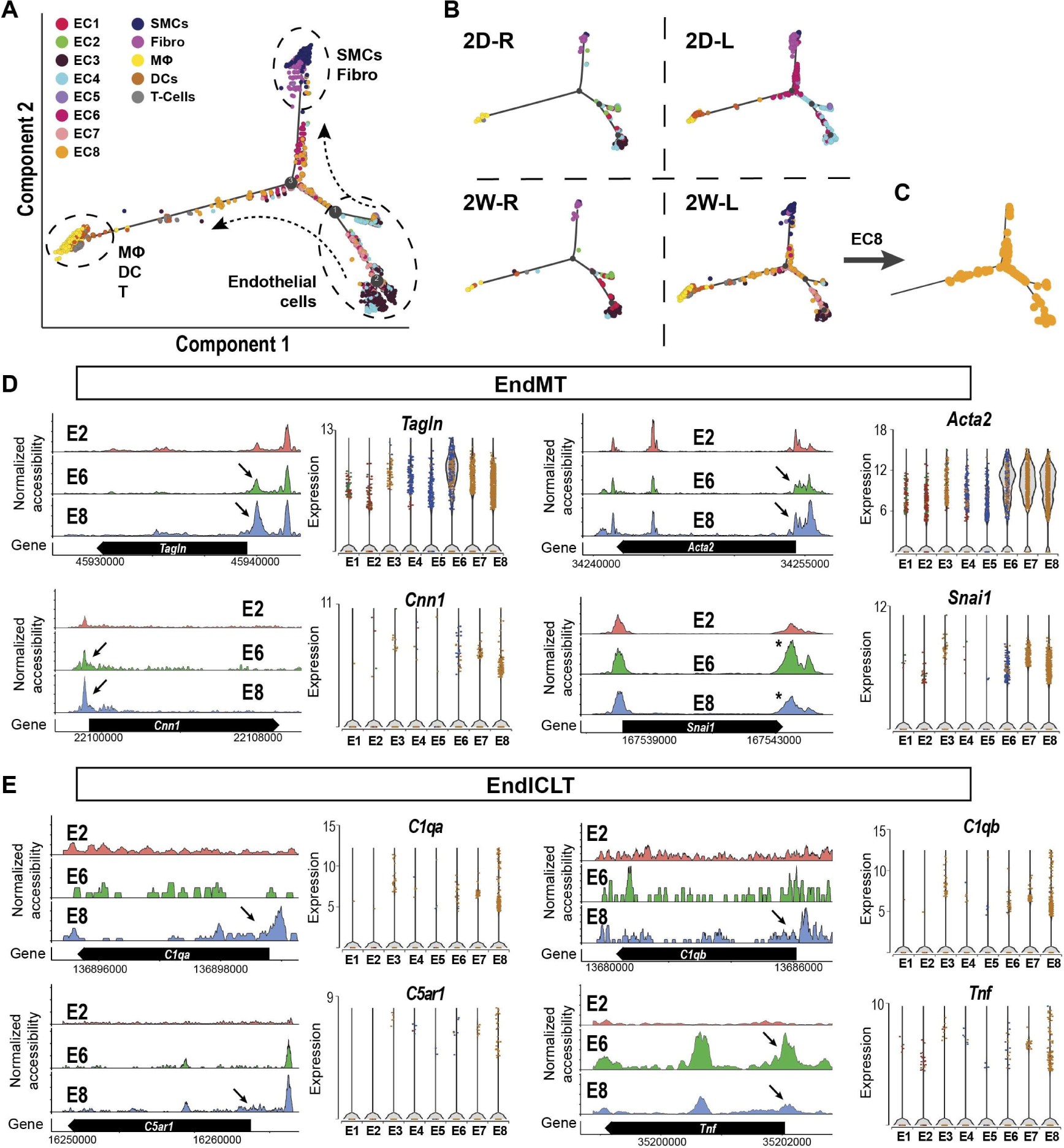
*D-flow* induced endothelial-to-mesenchymal transition (EndMT) and endothelial-to-immune like cell transition (EndICLT) as shown by psuedotime trajectory and accessibility profile analyses. **(A,B)** Pseudotime trajectory plot shows differentiated cell types (ECs, SMCs, Fibro and Mϕs, DCs and T-cells) at the end of the branches. Dots along the trajectory lines represent the status of the cells transitioning towards differentiated cell types. **(A)** Shows all cells pooled in 4 different conditions while **(B)** shows the cells split according to the time- and flow-dependent conditions. **(C and D)** The accessibility profiles of E1 and E2, representing ECs under *s-flow* and E8 representing ECs under *d-flow* from scATACseq data and violin plots showing the expression of corresponding genes from scRNAseq data are shown. Genes representing **(C)** and **(D)** EndICLT in the EC clusters are shown. Black arrows indicate accessibility changes in the promoter region except for *Snai1 (*)*.

To validate the transition of ECs to other cell types, EndMT and EndICLT, we further examined the chromatin accessibility and gene expression profiles of key marker genes. Figure 4D shows four markers of EndMT, *Tagln, Acta2, Cnn1*, and *Snai1* in EC clusters representing *s-flow* (E2), acute *d-flow* (E6), and chronic *d-flow* (E8) for their chromatin accessibility changes. *D-flow* increased the chromatin accessibility of these marker genes in the promoter regions (black arrows) or 3’UTR (see *). Consistent with the accessibility changes, the corresponding gene expression levels were also increased in ECs exposed to *d-flow* in E8. These results confirm that *d-flow* indeed induced EndMT as expected (Glaser et al., 2020; Lai et al., 2018; Mahmoud et al., 2017; Moonen et al., 2015; Ten Dijke et al., 2012; Warboys et al., 2019).

Having validated the EndMT, we further analyzed the gene markers of potential EndICLT using Mϕ markers, *C1qa, C1qb, C5ar1*, and *Tnf* in EC clusters. As shown in Figure 4D, chronic *d-flow* increased accessibility of these genes in their promoter regions (arrows) in E8 compared to the *s-flow* (E2) and acute *d-flow* (E6). This result was further confirmed by the increased expression of the corresponding genes in E8 as shown in the violin plot of the scRNAseq data (Figure 4E). These results further support that chronic exposure to *d-flow* can induce ECs transition toward immune-cell like populations, although not fully.

The EndMT and EndICLT results suggested that *d-flow* may induce broad endothelial reprogramming. To test the hypothesis, we examined the expression of marker genes for endothelial-to-hematopoietic transition (EndHT), endothelial stem cell (ESC), endothelial progenitor cell (EPC), and antigen-presenting cells transition (APC) (Santambrogio et al., 2019) using both the scRNAseq and scATACseq data sets. Table S3 shows that *d-flow* indeed induced expression of those marker genes of EndHT (*Sox7, Sox17, Gata2, Kit, Notch1, Eprc, Tie2, Bmp4*) and APCs (*Cd74, H2-aa, H2-ab1, H2-eb1*), as well as, previously explained EndMT (*Tagln, Cnn1, Acta2, Snai1)* and EndICLT (*C1qa, C1qb, C5ar1, Tnf, and Lyz2)*. Further, we found ECs expressing the marker genes of ESCs and EPCs in a flow-sensitive manner. ECs in E2 in *s-flow* were *Cd157*^high^*Sca1*^+^*Kit*^*-*^, E5/E6 in acute d-flow showed Cd157^low^*Sca1*^Low^*Kit*^-^, whereas E8 in chronic d-flow was *Cd157*^High^*Sca1*^Low^*Kit*^+^. We noted that ESCs were present in both E2 and E8, while EPCs were found only in E8. The underlying mechanisms, roles, and implications for these flow-sensitive changes in ESCs and EPCs need further investigation. Together, these results clearly demonstrate that *d-flow* reprograms ECs through EndMT, and potentially via EndICLT and EndHT by altering the gene expression of key genes at the transcriptional and chromatin accessibility levels.

We next analyzed the proatherogenic pathways activated by *d-flow* conditions identified from our gene ontology conditions (Figure 3B). As shown in Figure 5, *d-flow* in E8 altered markers of leukocyte traffic and inflammation (*Il1β, Il6, Ccl2*, and *Ccl3); ECM regulation (Adamts4, Lamb1, Timp3*, and *Timp4); vasoregulation (Nos3, Ptgs2*, and *Edn1);* and *lipid metabolism (Tm6sf2, Lsr* and *Mgll)*, in comparison to *s-flow* in E2 (Figure 5A-D). These results show that *d-flow* activated pro-atherogenic endothelial responses, while inhibiting anti-atherogenic pathways by changing the gene expression at the transcriptional and chromatin accessibility levels.

**Figure 5.**
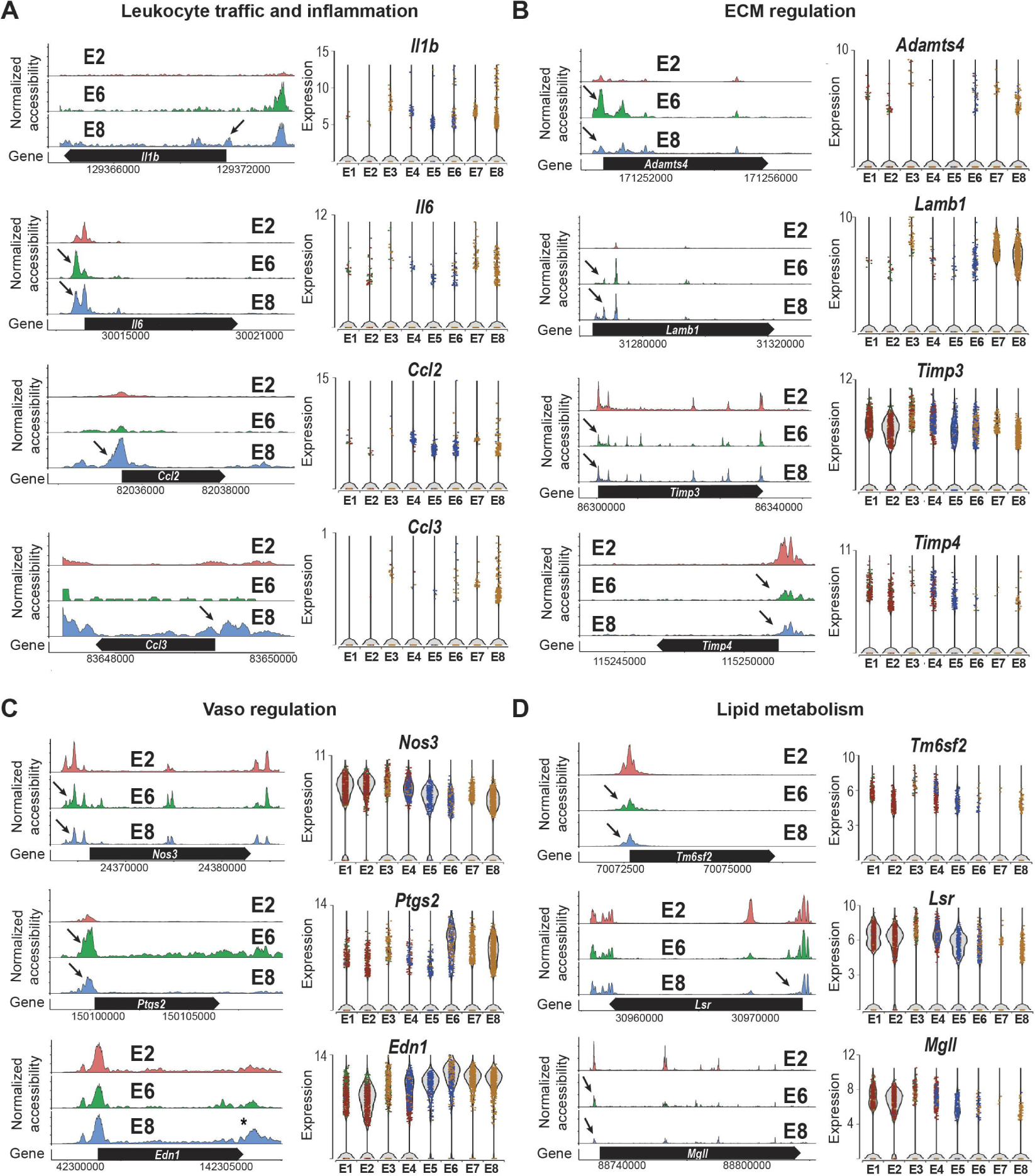
Chronic *d-flow* reprograms endothelial functions. The accessibility profiles of E1 and E2 (*s-flow)* and E8 (*d-flow)* from scATACseq data and violin plots showing the expression of corresponding genes from scRNAseq data are shown. Genes representing **(A)** leukocyte traffic and inflammation and **(B)** ECM regulation, **(C)** Vasoregulation and **(D)** Lipid metabolism in the EC clusters are shown. Black arrows indicate accessibility changes in the promoter region except for *Edn1 (*)*.

### Identification of novel flow-dependent TF binding motifs in vivo

We next examined if flow regulates gene expression through a set of TF binding motifs in response to either *s-flow* or *d-flow*. To test this hypothesis, we used Signac and ChromVar packages to identify flow-sensitive TF binding motifs using our scATACseq dataset (Figure 6). Top 20 TF binding motifs were identified in each endothelial cluster (E1-E8) and were plotted as a heatmap (Figure 6A). Figure 6B and C show TF binding motifs and the respective cell clusters where they were found. The TF binding motifs for KLF4 were enriched in E2 exposed to *s-flow*, while NRF1 was enriched in E3 and E4 (exposed to all flow conditions) (Figure 6B). In contrast, TF motifs for TEF, ETV3, RELA, FOS::JUN, TEAD1, and STAT1 were enriched in E5-E8 exposed to *d-flow* conditions (Figure 6C). Identification of the motifs for KLF4 (identical to the KLF2 motif), FOS:JUN (AP1), RELA, and TEAD1 as the most flow-sensitive TF motifs were expected, confirming the validity of our analysis (Bondareva et al., 2019; Fraineau et al., 2015; Kempe et al., 2005; Sangwung et al., 2017). We attempted to look for the shear stress response element (SSRE) (Resnick et al., 1993), but failed due to the lack of a defined list of TFs for the SSRE in our database. Our study validated the role of KLF2/4 as the master TF for *s-flow*, while identifying both known and novel TFs mediating the *d-flow* effects.

**Figure 6.**
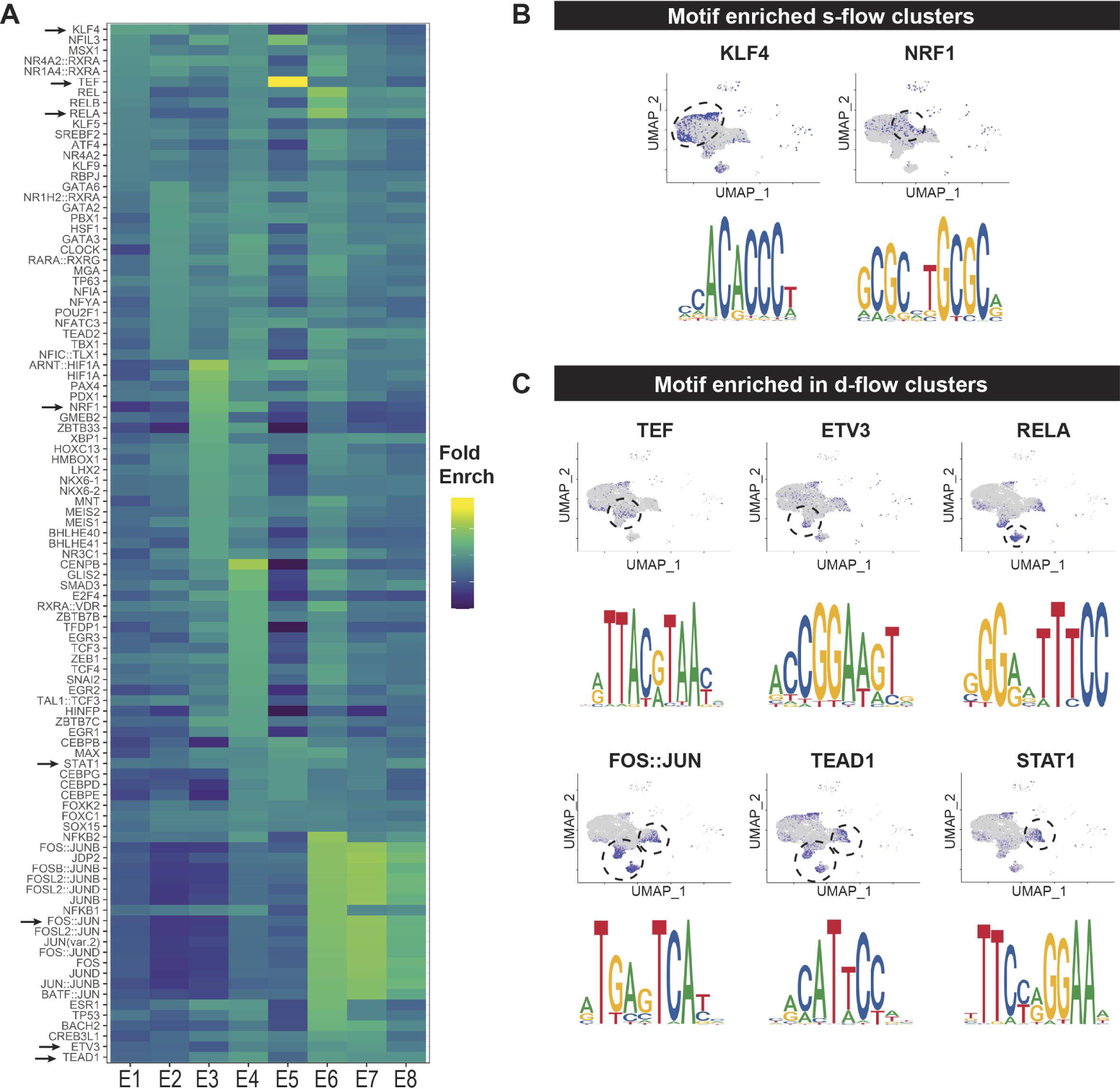
*D-flow* alters accessibility of transcription factors as determined by motif enrichment analysis. **(A)** Heatmap shows the top overrepresented motifs enriched in each endothelial cluster. Fold enrichment of each motif is represented on a color scale, dark blue being the lowest and yellow being the highest score. UMAP plot and the enriched motif sequences for overrepresented transcription factor binding motifs enriched in 2D-R and 2W-R (*s-flow*) **(B)**, and 2D-L (acute *d-flow)* and 2W-L (chronic *d-flow)* conditions **(C)**. The blue dots on the UMAP plot represent the cells where the motifs are overrepresented.

### Accessibility and co-accessibility analyses reveal flow-dependent cis-regulatory interactions

To identify flow-dependent regulation of cis-regulatory elements, enhancer regions, and co-accessibility changes, we carried out a Cicero analysis using our scATACseq data set. First, the prototypical mechanosensitive *Klf4* was analyzed, and the results were mapped to the corresponding gene locus using the ENCODE Genome Browser (Figure 7A). For this analysis, we compared E2 (*s-flow*) and E8 (chronic *d-flow*) clusters. E2 cells showed a higher co-accessibility between the cis-elements compared to E8, suggesting that *d-flow* decreased the co-accessibility connections. Next, as additional examples, we showed that the *d-flow* dramatically reduced the accessibility of the promoter region of *Arl4d* (Figure 7B-b). Similarly, the *d-flow* also reduced the co-accessibility and gene expression of *Arl4d* (Figure 7B-c and d). In contrast, the *d-flow* increased the accessibility, co-accessibility, and gene expression of *Tgfbi* (Figure 7C). These findings suggest that flow regulates the gene expression by regulating not only the TF binding motifs, but also by altering cis-regulatory elements and their interactions suggesting both epigenetic and transcriptional mechanisms.

**Figure 7.**
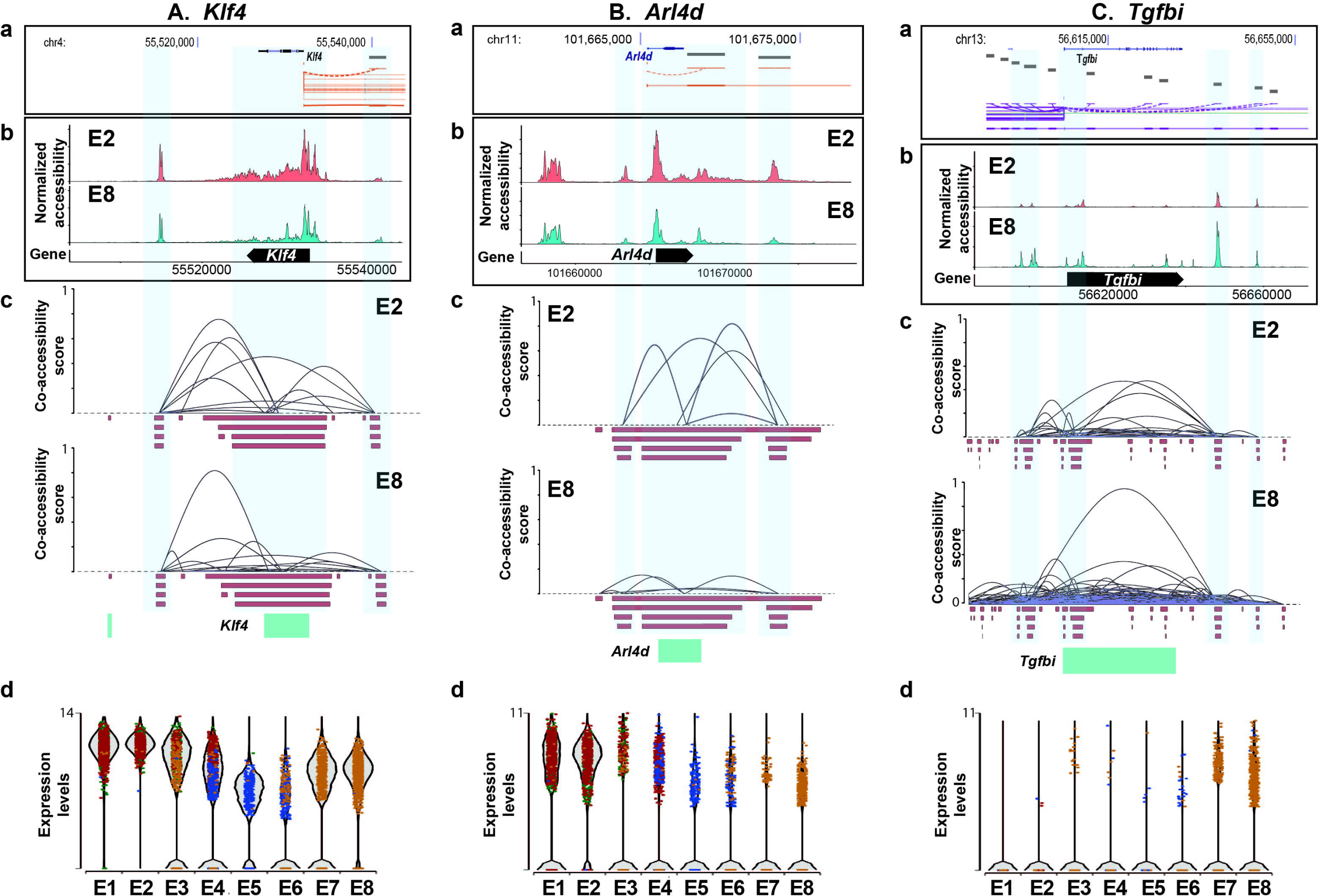
Chronic *d-flow* alters cis-regulatory interactions of flow-sensitive genes. Bioinformatic analysis showing the regulatory elements of **(A)** *Klf4*, **(B)** *Arl4d* and **(C)** *Tgfbi* genes. Gene structure and location of genes from ENCODE database **(a)** were mapped with normalized accessibility **(b)** and co-accessibility score plots **(c)** of E2 and E8 from our scATACseq dataset. **(d)** shows violin plots showing the expression of *Klf4* and *Arl4d* in E1 to E8 from our scRNAseq data.

## DISCUSSION

Here, we analyzed scRNAseq and scATACseq data sets and identified that 1) ECs are heterogeneous even under *s-flow* condition, 2) ECs are highly plastic, 3) *d-flow* dramatically alters endothelial phenotypes indicating that ECs transition from the quiescent, anti-inflammatory and anti-atherogenic phenotype to pro-inflammatory, pro-atherogenic phenotype, and 4) d-flow induces gene changes indicating that ECs undergo EndMT, EndHT, and EndICLT, while its effect on ESC is not clearly defined. In addition to 8 EC clusters, we identified SMCs, Fibro, Mϕs, DCs, and T-cells in our endothelial-enriched single cell populations obtained from the luminal flushings of the LCAs and RCAs. The comparison of the prototypical EC profiles (E8 by *d-flow* vs. E2 by *s-flow*) indicated that *d-flow* activated proatherogenic pathways including inflammation, leukocyte traffic, EndMT, ECM regulation, lipid metabolism, and vasoregulatory gene expression profiles. We also found TF binding motifs enriched by *s-flow* (KLF4/KLF2) and *d-flow* (TEF, RELA, ETV3, FOS::JUN, TEAD1, STAT1). Our study also showed that flow regulates chromatin co-accessibility and interactions among cis-regulatory elements. These results demonstrate that *d-flow* regulates gene expression both at the transcriptional and epigenetic levels, leading to the reprogramming of ECs into a mesenchymal, inflammatory, and immune cell-like phenotype.

Our scRNAseq and scATACseq results are consistent with our previous gene array results using the endothelial-enriched RNA samples in bulk obtained from the same mouse PCL model (Ni et al., 2010). As shown in our bulk RNA data and other reports (Ajami et al., 2017; Jiang et al., 2014; Lee et al., 2017; Ni et al., 2010; Son et al., 2013; Wang et al., 2016; Wang et al., 2006), our single-cell data showed expected changes in many well-known flow sensitive genes including *Klf2, Klf4, Nos3, Klk10, Angpt2, Ctgf, Bmp4, Timp3, Tgfb1, Thsp1, Col1a1, Tagln, Acta2, Cnn1*, and *Snai1*. The current study also showed that *d-flow* induced inflammatory and proliferative pathways, consistent with our bulk RNA array result (Ni et al., 2010), further demonstrating the validity of our single-cell results. One of the major limitations of the bulk RNA study was the inability to ascertain the cell-types where the gene changes were occurring. The endothelial-enriched RNA preparation method used for the bulk RNA array study included other cell types such as Mϕs and SMCs present in the carotid intima. While our previous bulk RNA results showed that *d-flow* increased overexpression of many pro-inflammatory genes and immune cell marker genes, it was unclear whether those changes were indeed contributed by ECs in response to *d-flow*. Current single-cell analysis has now addressed this limitation. We also compared our carotid endothelial gene expression data to the previously published endothelial heterogeneity data from murine ECs from various tissue types and arteries (Kalluri et al., 2019; Kalucka et al., 2020). While we found that most of the highly enriched genes (*Fbln5, Mecom, Gja4, Azin1, Sox17, Mgp, Nrp1, Kcnj8, Cytl1, Bmx, Eln, Fbln2, Gkn3*) found in the arteries were also present in all of our EC clusters showing no flow dependency. However, *Sema3g* was mostly found in our ECs exposed to s-flow (E1 and E2 cluster). Interestingly, none of the tissue-specific endothelial genes highly enriched in all the tissue types studied, including the brain, heart, lungs, and kidney were expressed at a very low level or not detected in our carotid artery EC clusters. This further highlights the endothelial heterogeneity and the impact of local cues on tissue-specific endothelial gene expression. Taken together, our single-cell study clearly demonstrates that *d-flow* induces expression of many pro-inflammatory, pro-EndMT, and pro-EndICLT genes, reprogramming ECs from atheroprotective phenotype under *s-flow* to pro-atherogenic phenotype under *d-flow* as discussed further below. Another unexpected finding was that *d-flow* induced EndHT, although its functional significance in atherosclerosis remains to be studied. Further, we found ESC markers in *s-flow* condition (E2: CD157^High^/Sca1^High^), which were transiently reduced by acute *d-flow* (E5/E6: CD157^Low^/Sca1^High^) but transitioning to a different condition by the chronic *d-flow* (E8: CD157^High^Sca1^Low^). Based on these complex changes in the marker genes, it is unclear whether *d-flow* alters the number of ESCs at present.

The cell clustering analyses using the scRNAseq and scATACseq data showed that ECs are heterogeneous within the carotid artery under the *s-flow* condition, which becomes even more diverse in response to *d-flow* conditions in a time-dependent manner. From our scRNAseq analysis, we found four EC clusters (E1-E4) under *s-flow* condition, two new populations in response to the acute *d-flow* (E5 and E7), and two additional EC populations (E6 and E8) under chronic *d-flow* condition. Likewise, scATACseq analysis identified four EC clusters (E1-E4) under *s-flow* condition, three new populations in response to the acute *d-flow* (E5-E7), and one additional EC population (E8) under chronic *d-flow* condition. Overall, the comparison of these two datasets and integrated analysis showed that most EC populations under *s-flow* and *d-flow* conditions were highly concordant, cross-validating each other’s data. However, in response to acute *d-flow*, there was a slight discordancy in the EC clusters and Mϕ clusters between the two datasets. While the underlying mechanism needs additional studies, we hypothesize that acute *d-flow* induced chromatin accessibility changes observed ECs in the scATACseq data preceded the gene expression changes by the scRNAseq data. Also, the gene activity index estimated from the scATACseq data is not as sensitive as the gene transcript data from the scRNAseq data, resulting in some discordancy. Wϕ clusters found in the scRNAseq as one cluster since we could not find discernable characteristics among them, and they largely overlapped with the single Mϕ cluster found from the scATACseq analysis. Taken together, our results demonstrate that *d-flow* induces a dramatic shift in endothelial phenotypes at the genomic and epigenomic levels, suggesting ECs transitioning from the athero-protective *s-flow* phenotypes to pro-atherogenic *d-flow* phenotypes.

The differential gene expression analysis of the scRNAseq data further demonstrated that flow reprograms ECs. We established that E2, E5, and E8 as representative endothelial phenotypes for atheroprotective *s-flow*, acute *d-flow*, and chronic proatherogenic *d-flow* conditions, respectively. E2 (*s-flow*) showed a high expression of *Klf2, Klf4*, and *Klk10*, well-known flow-sensitive genes (Ni et al., 2010; Sangwung et al., 2017), while E5 (acute *d-flow*) showed a high expression of *d-flow* induced proatherogenic genes (*Ctgf, Serpine1*, and *Edn1*) (Brooks et al., 2002; Wang et al., 2016). Interestingly, E8 (chronic *d-flow*) showed a high expression of not only the flow-dependent genes, but also markers of mesenchymal cells, hematopoietic cells, APCs and immune cells, indicating EndMT, EndHT, and the novel EndICLT (Figure 4 and Table S3). While *d-flow* is well-known to induce EndMT (Mahmoud et al., 2017; Moonen et al., 2015), it was unknown if *d-flow* can drive ECs to undergo EndHT and EndICLT status.

To further investigate whether EndICLT indeed occurs, we carried out the pseudotime trajectory analysis and found that chronic *d-flow* induced ECs not only to undergo EndMT as expected but also led to a transition towards immune cell-like phenotypes (Figure 4 and Figure S4). The EndICLT hypothesis was further validated by examining the expression of several gene markers of Mϕs (C*1qa, C1qb, C5ar1, Tnf and Lyz2*) by two independent analyses, scRNAseq and scATACseq (Figure 4D). These results demonstrate the novel role of *d-flow* in inducing EndICLT *in vivo*. The pseudotime trajectory analysis further reveals that *d-flow* drives ECs to either EndMT or EndICLT, but the underlying mechanism which determines these different EC fates is unknown. Our finding for EndICLT is supported by previous reports; vascular ECs and lymphatic ECs were shown to serve as APCs expressing MHC-I and MHC-II (Al-Soudi et al., 2017; Mai et al., 2013). The plasticity of ECs was further demonstrated by the transition of ECs to osteoblasts and hematopoietic cells (Guibentif et al., 2017; Lin et al., 2017). Interestingly, epithelial cell to immune cell-like transition was also reported (Choi et al., 2012). Moreover, recent studies comparing the translatomes and transcriptomes of various tissue-specific ECs demonstrated that are heterogeneous and plastic, adapting to the tissue-specific environment (Jambusaria et al., 2020; Kalucka et al., 2020). While the functional importance of EndMT in atherosclerosis is well established, the role of flow-dependent EndHT, EndICLT, and changes in ESC and EPC status in atherosclerosis will be exciting future research avenues.

Our motif analysis using the scATACseq data identified a comprehensive list of flow-sensitive TF biding motifs specific to *s-flow* and *d-flow*, respectively. As expected, KLF4 binding motifs were specifically enriched in ECs under *s-flow* condition. It is important to note that both KLF4 and KLF2 bind to the same motifs (Nayak et al., 2011). We identified several novel and expected TF binding motifs enriched under *d-flow* conditions. RELA, FOS::JUN and TEAD1 were previously reported as flow-sensitive TF binding motifs, while TEF, ETV3 and STAT1 as novel *d-flow* sensitive TF binding motifs (Figure 6). Our results validate the predominant role of KLK2/4-mediated gene regulation under *s-flow*, whereas *d-flow* regulates gene expression via several TFs. These results demonstrate that flow regulates access to these TF binding motifs via epigenomic mechanisms at a genome-wide level.

Our results should be interpreted with some caveats since the single cells and nuclei were obtained from the collagenase digestion, which could have affected the transcriptomic and epigenomic profiles during the incubation period (∼45 min). However, overwhelming list of flow-sensitive marker genes such as *Klf2, Klf4, Klk10*, and EndMT genes showed the robust expected flow-dependency, although we found some exceptions such as *Vcam1*.

In summary, our scRNAseq and scATACseq studies provide the comprehensive atlas of transcriptomic and epigenomic chromatin accessibility profiles at a single-cell level in response to *s-flow* and *d-flow in vivo*. We showed that flow alters the accessibility of TF binding motifs and cis-regulatory elements, thereby regulating gene expression at the genomic and epigenomic levels in ECs. These flow-dependent genomic and epigenomic changes resulted in endothelial heterogeneity and reprogramming, inducing EndMT, EndHT, and EndICLT. These robust and broad flow-dependent changes in ECs, from the quiescent and proatherogenic phenotype under *s-flow* condition to the pro-inflammatory and pro-atherogenic phenotype under *d-flow* condition, would lead to atherosclerosis in the presence of additional risk factors such as hypercholesterolemia. Our single-cell results would serve as valuable resources for the researchers in the field and harnessing the insights provided by our data would lead to a better understanding of flow-dependent regulation of endothelial biology and pathobiology. These insights will further facilitate the development of new therapeutics for atherosclerosis.

## ACKNOWLEDMENTS

This work was supported by funding from the National Institutes of Health grants HL119798, HL095070, and HL095070 to HJ. HJ was also supported by Wallace H. Coulter Distinguished Faculty Chair Professorship.

## AUTHOR CONTRIBUTIONS

Conceptualization, AA, SK, JK, and HJ; Investigation, AA, SK, JK, DK, HM, and JP.; Writing, AA, SK, JK, and HJ.; Funding Acquisition, HJ; Supervision, SK, HJ.

## STAR METHODS

### Mouse partial carotid ligation (PCL) surgery

All animal procedures used for this study were approved by the Institutional Animal Care and Use Committee at Emory University. To minimize sex-dependent variation and the complication of hypercholesterolemic conditions, we used a total of 44 male C57BL/6 mice (Jackson Laboratory, Bar Harbor, 10-week-old) which were fed *ad libitum* with standard mouse chow diet. To induce *d-flow*, PCL surgery was performed (Nam et al., 2010). Briefly, mice were initially anesthetized with 3.5% isoflurane and subsequently maintained on 1.5% isoflurane during the entire procedure. The LCA bifurcation was exposed by blunt dissection and three of four caudal LCA branches (left external carotid, internal carotid and occipital arteries) were ligated with 6-0 silk sutures, leaving the superior thyroid artery intact. The contralateral RCA was left untouched and served as an internal control. Following surgery, analgesic buprenorphine (0.1 mg kg^-1^) was administrated. At 2 days (2D-R and 2D-L) and 2 week post-partial ligation (2W-R and 2W-L), LCAs and RCAs were dissected out and a luminal enzymatic digestion of the carotids was performed to isolate endothelial-enriched single-cell preparations (Figure S1). Single-cell preparations from LCAs and RCAs from 12 mice were pooled to prepare single-nuclei for scATACseq study, while LCAs and RCAs from 10 additional mice were pooled to obtain single cells for scRNAseq study. Both studies were performed back to back.

### Single-cells and single-nuclei isolation and library preparation for scRNAseq and scATACseq analyses

After euthanizing the mice using CO2, the LCAs and RCAs were dissected, cleaned, and perfused with normal saline solution before a dissociation buffer was injected. The dissociation buffer was slightly modified from a previously reported recipe (Kalluri et al., 2019) by including 600 U ml^-1^ Type II Collagenase (02100502-CF, MP biomedicals, Irvine, CA) and 60 U ml^-1^ DNase I (M0303S, New England Biolabs Inc, Ipswich, MA) in 0.5 % fetal bovine serum (FBS) of 1X phosphate-buffered saline (PBS). Immediately after the introduction of dissociation buffer into the carotid lumens, the ends of the RCAs and LCAs were clamped and dissected out. The explanted carotids were incubated in 35-mm dishes containing HEPES-buffered saline solution (HBSS) at 37°C. After 45 minutes, the LCA and RCA lumens were flushed with the dissociation buffer into a 1.5 ml Eppendorf tube. The flushings were washed with ice-cold PBS and centrifuged for 5 minutes at 500 x g and were further incubated with accutase (A6964, Sigma-Aldrich, St. Louis, MO) for 5 minutes at 37 °C to prepare single-cells.

For scRNAseq, single-cells were resuspended in 1% BSA in PBS, counted and immediately processed at the Emory Integrated Genomics Core (EIGC) using a 10x Genomics Chromium device for single-cell encapsulation, barcoding, and RNA preparation. Subsequently, the cDNA libraries were prepared and sequenced on Illumina NextSeq® to a depth of 15,000 unique molecule identifier (UMI) per cell. UMI counts for each cellular barcode were quantified and used to estimate the number of cells successfully captured and sequenced.

For scATACseq, single-cells were washed with 0.04% BSA and incubated in a lysis buffer containing 10 mM Tris-HCl, 10 mM NaCl, 3 mM MgCl2, 0.1% Tween-20, 0.1% NP40, 0.01% digitonin and 1% BSA in nuclease-free Water. After 5 min, the lysates were washed with wash buffer (10 mM Tris-HCl, 10 mM NaCl, 3 mM MgCl2, and 0.1% Tween-20) and the nuclei were resuspended in a Nuclei Buffer® (10x Genomics), and were immediately processed at the EIGC for scATAC sequencing.

The nuclei (∼7,000 each) from 2D (R and L) and 2W (R and L) were individually incubated with ATAC Buffer (10x Genomics) and ATAC Enzyme (Tn5 transposase, 10x Genomics) to form a transposition mix for 60 min at 37°C following manufacturer’s protocol. A mild detergent condition was chosen to keep the nuclei intact during tagmentation. Then a master mix composed of Barcoding Reagent, Reducing Agent B and Barcoding Enzyme (10x Genomics) was added. The resulting mixtures were loaded onto a Chromium Chip (10x Genomics; 2000121) and single-nuclei gel emulsions with barcoding were prepared following the manufacturer’s instructions (10X Genomics).

The Cell Ranger Single-Cell Software suite was then used for demultiplexing, barcode processing, alignment, and initial clustering of the raw scATACseq and scRNAseq profiles.

### scRNAseq data analysis

We analyzed the scRNAseq data with Seurat R package. Briefly, the scRNAseq data files were processed with Cell Ranger Software (version 3.1; https://support.10xgenomics.com/single-cell-atac/software/pipelines/latest/algorithms/overview) where the sequencing reads were aligned to mouse reference genome using STAR aligner. A read was considered exonic, if at least 50% of it mapped to an exon, intronic (if it was non-exonic and intersected an intron), or intergenic otherwise. For reads that aligned to a single exonic locus but also aligned to 1 or more non-exonic loci, the exonic locus was prioritized and the read was considered to be confidently mapped to the exonic locus. Cell Ranger also aligned exonic reads to annotated transcripts. An annotated transcript that aligned to the same strand was considered to be confidently mapped to the transcriptome. These confidently mapped reads were used for unique molecular identifier (UMI) counting and subsequent analysis to generate h5 files. The h5 file of each sample was then processed with Seurat R package for further analysis. First, a quality check was performed to remove low-quality cells and doublets by selecting the cells containing unique feature counts of over 200 or less than 7,600, and cells with lower than 10% of mitochondrial counts. All data sets were merged, normalized, scaled, clustered, and visualized by UMAP.

As an independent validation, the scRNAseq analysis was also carried using Partek Flow® analysis software following the same filtering criteria. Both Partek Flow and Seurat analyses provided consistent and nearly identical results.

### Single-cell ATACseq data analysis

The sequencing data for the scATAC study were processed by Cell Ranger (CR) Software (version 1.2; https://support.10xgenomics.com/single-cell-atac/software/pipelines/latest/algorithms/overview) as described (Satpathy et al., 2019). Briefly, to avoid barcode misidentification, each 16-nucleotide barcode was compared against a ‘whitelist’ of correct barcode sequences, and the frequency of each whitelist barcode is counted. The barcodes with less than 2 nucleotide mismatches were corrected and included for further analysis. Prior to the alignment of the sequences to a mouse reference sequence (mm10), the *cutadapt* tool was used to trim the barcodes using default parameters. Reads with less than 25 bp were not included in further analysis. Fragments were then identified as reading pairs if their MAPping Quality (MAPQ) scores were larger than 30 on both reads, not mitochondrial, and not chimerically mapped. Once the fragments were filtered, fragments.tsv.gz file was created marking the start and end of the fragment after adjusting the 5’ ends of the read-pair to account for transposition. The file was position-sorted and ran through the SAMtools tabix command with default parameters. Peak calling was also performed with Cell Ranger using the sites as determined by the ends of the fragments in the position-sorted fragments.tsv.gz file, which counts the number of transposition events at each base-pair along the genome. A signal threshold was set up based on an odds-ratio of 1/5 that determines whether a region is a peak signal or noise. Consequently, not all cut sites were within a peak region. Peaks within 500 bp of each other were merged to produce a position-sorted BED file of peaks. Finally, the Peak-Barcode Matrix was produced which consisted of the counts of fragment ends within each peak region for each barcode. This raw Peak-Barcode Matrix captures the enrichment of open chromatin accessibility per barcode.

The h5 files generated in Cell Ranger were subsequently analyzed using Signac which is an extension of Seurat R package for the analysis, interpretation, and exploration of single-cell chromatin datasets. First, for each sample dataset, (2D-R, 2D-L, 2W-R and 2W-L), a Seurat object was created, which contained the Peak Barcode-Matrix and the fragments file. Once the data objects were generated, the enrichment of Tn5 integration events at transcriptional start sites (TSS) was calculated for each sample. As a quality control, the cells with TSS enrichment score >1 and cells with at least 500 peaks were retained and used for further analysis.

The filtered datasets were then merged and the frequency-inverse document frequency (TF-IDF) normalization was performed followed by feature selection and dimensional reduction which returned a low dimensional representation of the object (analogous to the principal component analysis used in scRNAseq). Next, a graph-based clustering and non-linear dimension reduction were performed for UMAP visualization.

The analysis of differentially accessible regions between different cell clusters was performed with Signac, which follows a logistic regression and takes into account the differential sequencing depth (Stuart et al., 2019) (https://satijalab.org/signac/articles/mouse_brain_vignette.html).

Furthermore, chromatin accessibility was studied by Monocle 2 and Cicero R packages through the pseudotime trajectory analysis from scATACseq data (https://cole-trapnell-lab.github.io/cicero-release/docs_m3/#single-cell-accessibility-trajectories). In addition, cis-regulatory networks information obtained using Cicero was applied to our scATACseq dataset following this vignette https://cole-trapnell-lab.github.io/cicero-release/docs/#constructing-cis-regulatory-networks (Pliner et al., 2018).

Moreover, motif analysis was performed with Signac by two complementary methods (https://satijalab.org/signac/articles/motif_vignette.html). The first one performed a hypergeometric analysis to test the probability of observing the motif at the given frequency by chance comparing with a background set of peaks matched for GC content. The second one identified a differential motif activity analysis using *ChromVar*. In both cases, default parameters were used.

### Integration of scRNAseq and scATACseq datasets

Once the scRNAseq and scATACseq datasets were individually analyzed using the same Seurat v 3.6 package, they were further integrated using the integration vignette as described by Satija laboratory (Stuart et al., 2019). Briefly, for scATACseq, the gene activity matrix was computed, followed by normalization, reduction of dimensions and UMAP plot. For scRNAseq, the analytical results generated using Seurat were used for integration with scATACseq data. Next, scRNAseq and scATACseq anchors were identified and the cell cluster labels identified from the scRNAseq analysis were used for integration of scATACseq dataset. Finally, to plot both datasets together, scRNAseq and scATACseq cells were co-embedded in the same low dimensional space, merged and visualized by UMAP.

### Gene ontology analysis and Functional pathway analysis

Gene ontology analysis was performed using PANTHER (Protein Analysis Through Evolutionary Relationships, http://www.pantherdb.org/). Moreover, the term used in these analyses was GO molecular function. For functional pathway analysis, expression data from both scRNAseq and scATACseq analyses were used to generate differentially expressed genes or accessible genes, respectively, among cell clusters identified by the UMAP analyses. A *p*-value of <0.05 and a false discovery rate (FDR) of <0.05 served as cutoffs for significance between the groups. Significantly up-regulated genes between a group and all remaining groups were selected as the exclusively regulated genes per group. These genes were used for further pathway analysis using Panther (Yuzwa et al., 2017). Significantly regulated functional clusters or single pathways were further grouped by the indicated functional classes and compared by the enrichment score. Significantly enriched (p<0.05) canonical pathways were selected as representative pathways for each cluster. Additionally, gene ontology and functional pathway analysis were further validated by using Partek Flow® software using internal modules following the above-mentioned parameters.

## Data and Code availability

The scRNAseq and scATACseq sequencing data is accessible from the NCBI sequence read archives under the BioProject accession number PRJNA646233. The basic scripts for scRNAseq and scATACseq analysis can be accessed here: https://github.com/JoLab-Emory/SingleCell. Additional relevant data are available from the corresponding author.

