## Supplementary Figures and Tables for "Endothelial reprogramming by disturbed flow revealed by single-cell RNA and chromatin accessibility study"

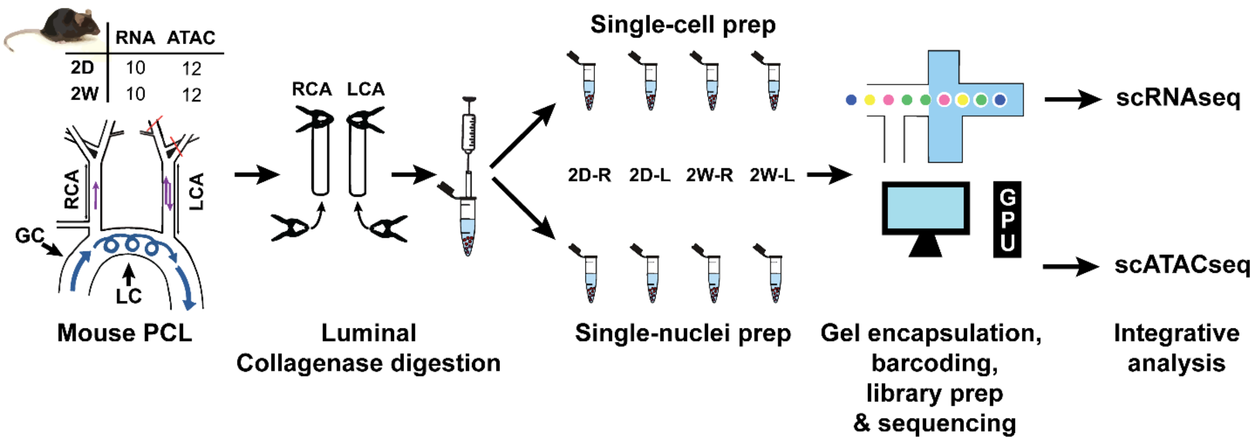


### Figure S1

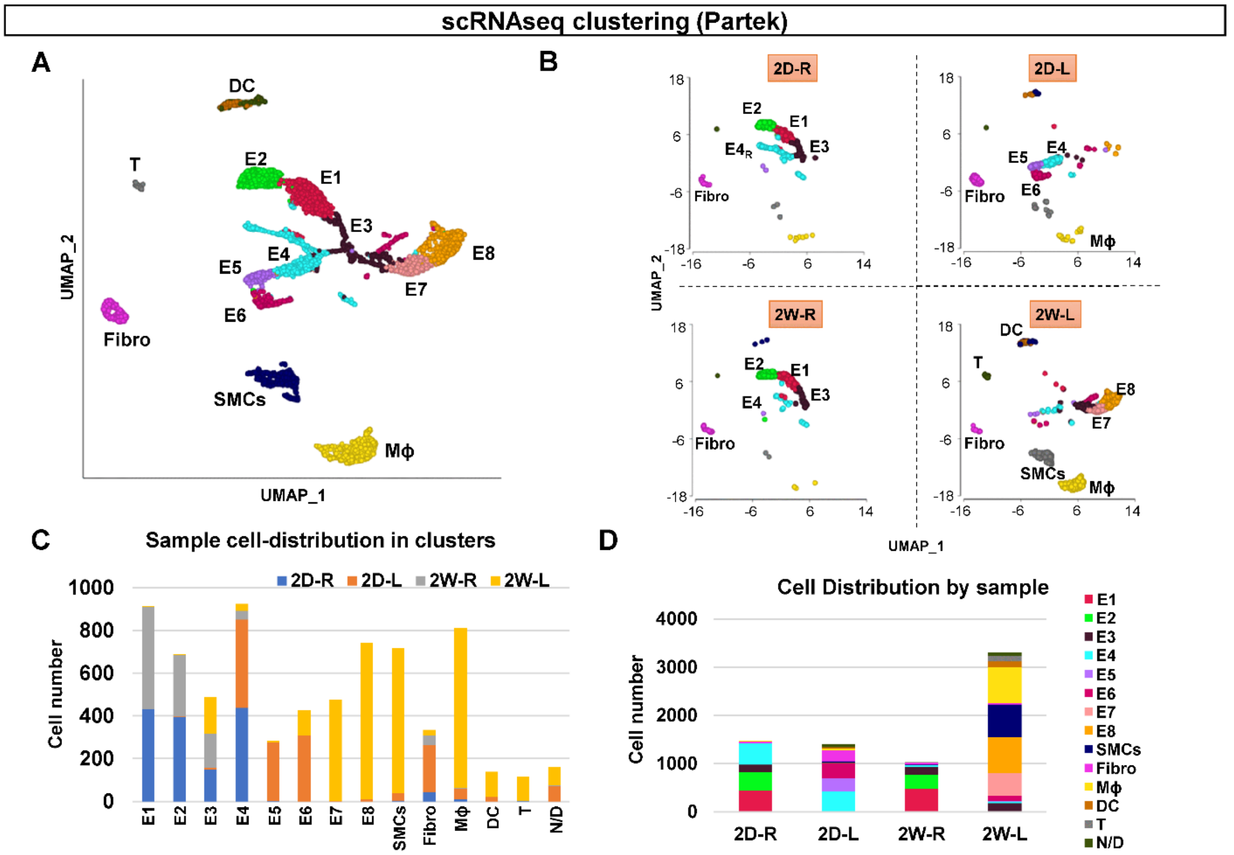


### Figure S2

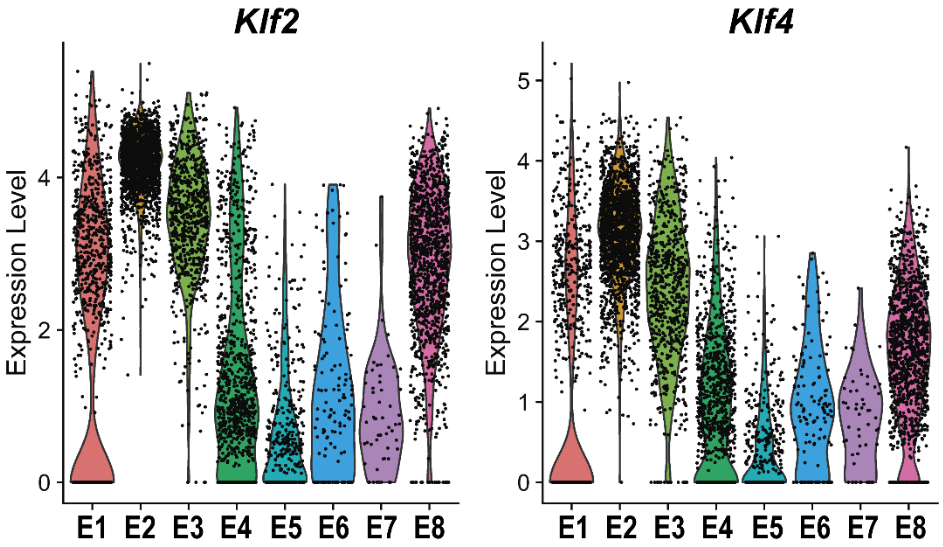


### Figure S3

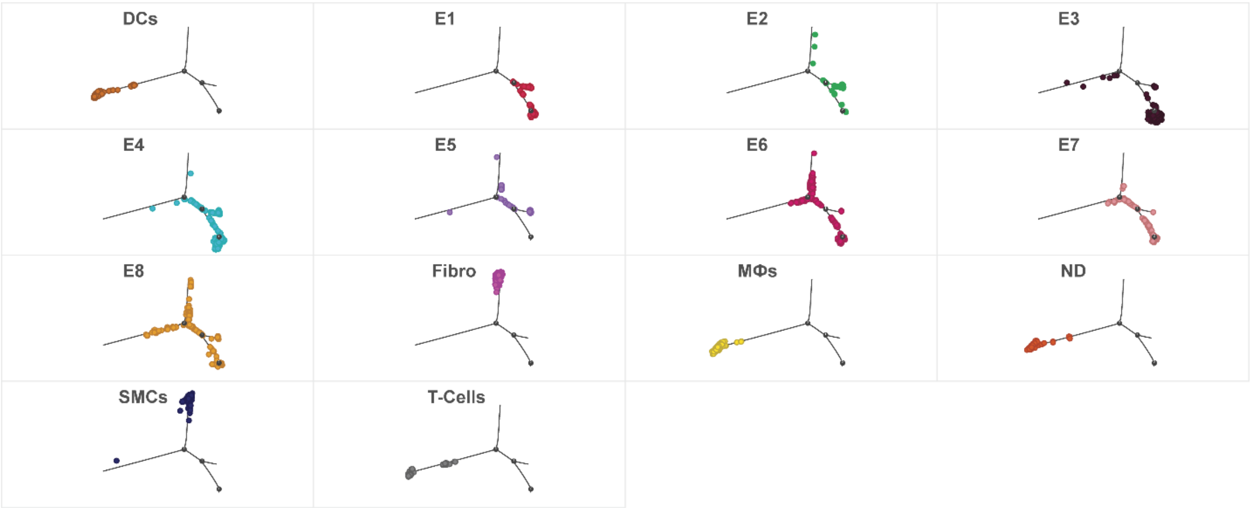


### Figure S4

### SUPPLEMENTARY FIGURES LEGENDS

**Figure S1. Overview of scRNA and scATAC studies.** Partial carotid ligation (PCL) was performed on mice (n=44) to induce *d-flow* in the left carotid arteries (LCA) using the right (RCA) as a *s-flow* control. Single cells were enzymatically isolated from the lumens of RCAs and LCAs at 2 days and 2 weeks post PCL. For scRNAseq, 10 RCAs and LCAs were pooled at each timepoint and single-cells were encapsulated, barcoded, cDNA library was prepared, and sequenced. For the scATACseq study, 12 RCAs and LCAs were pooled at each time point to prepare single-nuclei, which were encapsulated, treated with transposase, barcoded and sequenced. scRNAseq and scATACseq data were analysed with Seurat, respectively.

**Figure S2. scRNAseq clustering and cell identification using Partek Flow® software. (A and B). (A and B)** scRNAseq data representing 2-day RCA and LCA (2D-R and 2D-L) and 2-weeks RCA and LCA (2W-R and 2W-L) were plotted on a single UMAP to identify major cell populations. **(A)** shows a UMAP representing all 4 samples while **(B)** represents individual UMAP plots for each condition. Major cell populations include endothelial cells (ECs, E1-8), smooth muscle cells (SMCs), fibroblasts (Fibro), Monocytes/Macrophages (M$\phi$), dendritic cells (DCs), and T-cells (T). **(C)** Graph shows the cell numbers in each cell cluster for 4 different experimental conditions whereas **(D)** represents the number of cells in each sample and cluster.

**Figure S3. D-flow reduces the gene expression of the master regulators, *Klf2* and *Klf4*, in endothelial clusters. (A and B)** scRNAseq based violin plot representing the gene expression levels of Klf2 and Klf4, respectively through E1 to E8.

**Figure S4. scRNAseq based pseudotime trajectory analysis divided by cluster.** Each cluster is colored and labeled with its corresponding color and name.

### SUPPLEMMENTARY TABLES

#### *Table S1. scRNAseq information*

| Sample | Mice # | Experim. | Single-cell | Mean reads/ cell | Genes/ cell |
|---|---|---|---|---|---|
| 2D-R | 10 | 2 day, RCA | 1,867 | 55,593 | 3,376 |
| 2D-L | | 2 day, LCA | 2,119 | 76,050 | 3,284 |
| 2W-R | | 2 wk, RCA | 1,263 | 88,631 | 3,537 |
| 2W-L | | 2 wk, LCA | 4,460 | 34,142 | 2,770 |

#### *Table S2. scATACseq information*

| Sample | Mice # | Experim. | Single-cell | Mean fragments / cell | Total Reads |
|---|---|---|---|---|---|
| 2D-R | 12 | 2 day, RCA | 1,291 | 24,459 | 304,487,130 |
| 2D-L | | 2 day, LCA | 5,351 | 22,201 | 372,162,757 |
| 2W-R | | 2 wk, RCA | 5,826 | 19,690 | 336,611,489 |
| 2W-L | | 2 wk, LCA | 5,856 | 22,398 | 370,557,062 |

#### *Table S3. Chronic d-flow induces genes related with EndMT, EndICLT, EndoHT, and APC.*

| ***Process*** | ***Genes names*** | ***s-flow (E2 cluster)*** 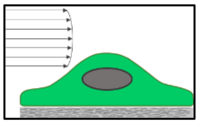 | ***Acute d-flow (scRNAseq: EC5 & scATACseq EC6)*** 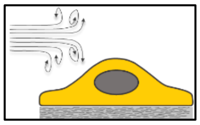 | ***Chronic d-flow (E8 cluster)*** 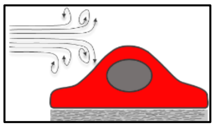 | ***References*** |
|---|---|---|---|---|---|
| ***Endothelial markers*** | *Pecam 1* | +++ | ++ | ++ | *(Kalluri et al., 2019)* |
| | *Cdh5* | +++ | +++ | +++ | |
| | *Icam2* | +++ | ++ | ++ | |
| | *Tie1* | +++ | ++ | ++ | |
| ***EndMT*** | *Tagln* | - | +++ | ++ | *(Lai et al., 2018; Mahmoud et al., 2017; Moonen et al., 2015)* |
| | *Cnn1* | -- | -- | + | |
| | *Acta2* | - | +++ | +++ | |
| | *Snai1* | -- | - | ++ | |
| ***EndHT*** | *Sox7* | +++ | + | + | *(Ottersbach, 2019)* |
| | *Sox17* | +++ | + | + | |
| | *Gata2* | +++ | + | + | |
| | *Kit* | -- | - | ++ | |
| | *Notch1* | +++ | + | + | |
| | *EPRC* | + | +++ | ++ | |
| | *Tie2* | +++ | + | + | |

| | | | | | |
|---|---|---|---|---|---|
| | *Bmp4* | + | ++ | +++ | |
| ***EndSC*** | *CD157* | ++ | - | ++ | *(Wakabayashi et al., 2018)* |
| | *Sca1* | ++ | | | *(Xiao et al., 2006)* |
| ***APC*** | *H2-aa* | - | - | ++ | *(Santambrogio et al., 2019)* |
| | *H2-Ab1* | - | - | ++ | |
| | *H2-Eb1* | - | - | ++ | |
| | *Cd74* | - | - | ++ | |
| ***EndICLT*** | *C1qa* | -- | -- | + | |
| | *C1qb* | -- | -- | + | |
| | *C5ar1* | -- | -- | + | |
| | *Tnf* | -- | -- | + | |
| | *Lyz2* | -- | -- | ++ | |
